# *OsJAZ9* overexpression improves potassium deficiency tolerance in rice by modulating jasmonic acid levels and signaling

**DOI:** 10.1101/440024

**Authors:** Ajit Pal Singh, Bipin K. Pandey, Poonam Mehra, Ravindra Kumar Chandan, Gopaljee Jha, Jitender Giri

**Author notes:** equal contribution (Ajit Pal Singh, Bipin K. Pandey). **Highlights:** OsJAZ9, modulates JA signaling and content in rice and modulates root system architecture and physiology of plant so that it can better tolerate K deficiency and sheath blight disease. Author for correspondence Jitender Giri 91-11-26735227.

## Abstract

Potassium (K) which makes around 2-10% of plants total dry biomass, when become deficient, makes the plants highly susceptible to both abiotic and biotic stresses. Recent evidences suggest overlapping transcriptional responses to K deficiency and Jasmonate (JA) treatment in plants. However, a link between these responses was missing. Notably, K deficiency and JA application produce similar phenotypic and transcriptional responses. Here, we used molecular, physiological and morphological studies to analyze the role of *OsJAZ9* in JA homeostasis, K deficiency and sheath blight resistance. We raised *OsJAZ9* overexpression, knockdown, translational reporter and C-terminal deleted translational reporter lines in rice to establish the role of JA signaling in K ion homeostasis and OsJAZ9 as a critical component of JA signaling for K deficiency response. *OsJAZ9* overexpression and knockdown provide K deficiency tolerance and sensitivity, respectively, by modulating various K transporters and root system architecture. Furthermore, RNA Seq and JA profiling revealed an elevation of JA responsive genes and JA levels in *OsJAZ9* OE lines under K deficiency. Our data provide clear evidence on the crucial role of JAZ repressor, OsJAZ9 in improving K deficiency tolerance in rice by altering JA levels and signaling.

## INTRODUCTION

Rice is a mainstay for global food security. Two third population of the world is highly dependent on rice as a staple food crop. Soil nutrient deficiency is one of the primary limitations to rice yield. Notably, Potassium (K) is one of the most important and abundant macronutrients in plants. It can comprise 2-10% of the plant’s total dry weight (Leigh & Wyn Jones, 1984). Its physiological functions fall into two categories; the first one requires high and relatively stable concentration of K^+^ to regulate the osmotic potential of the cell and activation of many enzymes involved in respiration and photosynthesis (K^+^ act as a cofactor for many enzymes). K^+^ also stabilizes protein synthesis and neutralizes the negative charges on proteins (Marschner 1995). The second category is based on the high mobility of K^+^ ions resulting into osmotic changes, which regulate the stomatal movement (Sawhney & Zelitch, 1969; Terry & Ulrich, 1973) and phloem transport (Mengel and Viro, 1974). K^+^ is also involved in various cellular processes such as energy production, cell expansion and balancing the counter flux of charges across the membranes (Tester and Blatt, 1989; Wu et al., 1991; Elumalai, 2002). Hence, its deficiency affects fitness and overall growth as K deficient plants are more susceptible to salt (Kaya et al., 2006), drought (Egilla et al., 2001; Wang et al., 2013), chilling (Kant and Kafkafi, 2002) and biotic stresses (Hardter, 2002; Sarwar, 2012). Therefore, its deficiency results in the highest crop yield loss, both directly and indirectly.

Although K is among the most abundant minerals in the soil; its availability to plants is limited because of the presence of K (about 98%) as minerals or non-exchangeable form where its release into the soil solution is far slower than the rate of acquisition by roots (Sparks, 1987). Notably, a very little fraction of soil K is present in the soil solution or exchangeable form whose availability again depends on multiple factors like soil pH (Rich and Black, 1964; Rich, 1964), presence of other monovalent cations like Na^+^ and NH_4_^+^ (Qi and Spalding, 2004) and the type of soil particles (Sparks, 1987).

K deficiency results in both root (Singh et al., 2015) and shoot (Shankar et al., 2013) growth inhibition. In roots, it strongly impairs primary root growth (Gruber et al., 2013); a similar phenotype observed on exogenous Jasmonic Acid (JA) application (Staswick et al., 1992; Cai et al., 2014). Further, few transcriptome studies have shown many JA signaling genes upregulated under K deficiency (Ma et al., 2012; Shankar et al., 2013; Takehisa et al., 2013). Interestingly, a substantial part of K-responsive transcriptome was either absent or replaced in Arabidopsis JA receptor, *coil-16* mutant (Armengaud et al., 2004; Ma et al., 2012; Takehisa et al., 2013) which indicates active roles of JA signaling in K deficiency responses. Moreover, a significant number of genes were found commonly upregulated by JA treatment and K deficiency in rice (Kobayashi et al., 2016), which suggest crosstalk between JA signaling and K deficiency response.

JA and its derivatives commonly called Jasmonates (JAs) are a group of oxylipin-derived phytohormones, which are synthesized in response to a large number of biotic and abiotic stresses experienced by plants (Feys et al., 1994; Kazan, 2015; Riemann et al., 2015). JAs are perceived simultaneously by a receptor/co-receptor complex called JAZ-COI1 complex (Sheard et al., 2010). COI1 (CORONATINE INSENSITIVE 1), a F-box E3 ubiquitin ligase, is co-receptor of JA and is a part of the ubiquitin ligase complex (Wasternack and Hause, 2013). JAZ (Jasmonate ZIM domain) proteins remain bound to JA responsive genes; bHLH transcription factors (TFs) and repress their activity. On the perception of JA-Isoleucine (JA-Ile, bioactive jasmonate form), COI1 targets JAZ protein for 26S proteasome-mediated degradation and hence, de-repress bHLH TFs allowing the expression of genes responsive to JAs (Wasternack and Hause, 2013). JAZ protein contains a highly conserved N-terminal ZIM domain (or TIFY motif) which is responsible for homo or hetero-dimerization of JAZ proteins and recruits NINJA (Novel interactor of JAZ)/TOPLESS for repressor activity (Chung et al., 2009; Pauwels et al., 2010), and a C-terminal Jas domain which binds with MYC2 for its transcriptional repression (Chini et al., 2007; Thines et al., 2007). Jas domain also facilitates the interaction of JAZ proteins and COI1; therefore, it also allows the degradation of JAZ protein in the presence of JA-Ile (Wasternack and Hause, 2013). JAZs form a small family of 15 members in rice (Ye et al., 2009). While their roles have been established in floral development, salt, and biotic stress tolerance, their functions in K deficiency remain elusive. Here, we determine that OsJAZ9 is an integral player of JA mediated K deficiency tolerance in rice. Overexpression of *OsJAZ9* enhanced both JA biosynthesis and signaling in rice, which also improved resistance against necrotrophic pathogen *Rhizoctonia. Solani.*

## Material and Methods

### Plant growth conditions

Rice (*Oryza sativa* var PB1) seeds were surface-sterilized with 0.1% HgCl_2_ for 10 min and then washed with sterile water five times. After that, seeds were germinated on autoclaved wet tissue paper for two days in the dark followed by two days in the light. Equally germinated seeds were then transferred to full strength liquid Yoshida media with NH_4_NO_3_ (1.40 mM), NaH_2_PO_4_ (0.32 mM), K_2_SO_4_ (0.51 mM), CaCl_2_.2H_2_O (1 mM), MgSO_4_.7H_2_O (1.7 mM), H_3_BO_3_ (19 μM), ZnSO_4_.7H_2_O (0.15 μM), CuSO_4_.5H_2_O (0.15 μM), (NH_4_)_6_Mo_4_O_2_.4H_2_O (0.015 μM), Citric Acid (70.75 μM), Na-Fe-EDTA (60 μM) and MnCl_2_.4H_2_O (9.46 μM). The concentration of K_2_SO_4_ was adjusted as required to make-up the desired concertation of K. All the experiments were carried out in a controlled growth chamber maintained at 16 h photoperiod, 30°C day and 28°C night temperature, 280-300 μM photons/m2/sec photon density and □ □ 70% relative humidity.

### qRT-PCR analysis

Gene-specific primers were designed from CDS (retrieved from Rice Gene Annotation Project) using PRIMER EXPRESS version 3.0 (PE Applied Biosystems TM, USA) with default parameters. SYBR® Green Master Mix was used to quantify the DNA product in Applied Biosystems 7500 Fast Real-Time PCR. For the Relative gene expression analysis, we used the ΔΔCt method taking *Ubiquitin5* (Os01g0328400) as an endogenous control (Mehra et al., 2016).

### Measurement of K and Na

Root and shoot tissues were harvested separately and repeatedly washed with milliQ water. Dried tissues were ground, and 50 mg sample. The diluted cooled digest was filtered and stored in polypropylene bottles with a final volume of 50 ml. Analysis of Na and K content was performed using a flame photometer (Systronics Flame Photometer 128μC).

### Generation of transgenic lines

Full-length cDNA (AK070649) of *OsJAZ9* was amplified using gene-specific primers (Supplemental Table S1) and cloned under *ZmUbi1* promoter in gateway-compatible overexpression vector as described (Pandey et al., 2017). For raising RNAi transgenics of *OsJAZ9*, 416 bp region of *OsJAZ9* cDNA was amplified from the cDNA region encompassing 3’UTR and cloned in pANIC8b vector (Mann et al., 2012) by Gateway Technology. To generate *p35S:OsJAZ9:GUS* and *p35S:OsJAZ9ΔC:GUS* lines, *OsJAZ9* coding region and *OsJAZ9*Δ*C* (deleted 65 amino acids including jas motif from the C-terminal of OsJAZ9) cDNAs were amplified using gene-specific primers, and cloned into pCAMBIA1301. Overexpression, RNAi, *p35S:OsJAZ9:GUS* and *p35S:OsJAZ9AC:GUS* transgenic rice lines were raised as described (Mehra et al., 2017). Regenerated plants were selected on hygromycin (50 μg/ml) and screened by qRT-PCR and GUS histochemical staining. All the experiments were performed in T3 homozygous transgenic lines.

### Analysis of GUS Activity

Newly emerged roots were selected for histochemical staining, i.e. gus activity. Samples were harvested at indicated time points and immersed in GUS buffer (1 mM 5-bromo-4-chloro-3-indolyl-β-glucuronide sodium salt in 50 mM sodium phosphate, pH 7.0, ten mM Na-EDTA, 0.5 mM ferricyanide, 0.5 mM ferrocyanide, and 0.1% Triton X-100), and were incubated at 37°C for overnight. Chlorophyll was cleared from plant tissues by immersing them in 70% ethanol, overnight. For detection of *OsJAZ9* degradation in-vivo, roots from transgenic lines containing *p35S:OsJAZ9:GUS* and *p35S:OsJAZ9ΔC:GUS* plants were treated with MeJA (100 pM) with or without the proteasome inhibitor MG132 (100 μM) for 1h. Histochemical GUS staining of roots was performed to visualize the abundance/degradation of OsJAZ9 and OsJAZ9AC. GUS signal was observed under a stereo-zoom microscope.

### Subcellular localization of OsJAZ9

CDS of OsJAZ9 was cloned into pSITE3CA vector to produce YFP:OsJAZ9 fusion protein. Subcellular localization was analyzed in onion epidermal cells using the particle bombardment method (Singh et al., 2015). YFP fluorescence was visualized under a fluorescence microscope (Nikon Eclipse 80i).

### MeJA mediated root growth inhibition assay

Uniformly, germinated seeds were transferred to Yoshida media and allowed to grow for ten days under normal conditions. After ten days of healthy growth, ten μM MeJA dissolved in DMSO was supplied to one set of plants (including WT, RN12, OE3, and OE10) while another set (control) was supplied with DMSO alone (mock). pH was maintained at 5.4 and media was replenished after five days. Root growth inhibition was analyzed after ten days of MeJA treatment by calculating the percentage change in root growth between DMSO and MeJA treated plants.

### Exogenous MeJA treatment to analyze the impact on K deficiency

To investigate the impact of exogenous application of MeJA on K deficiency, uniformly germinated WT rice seedlings were transferred to hydroponic (Yoshida) media with 4.08 μM K_2_SO_4_, 8.16 μM K_2_SO_4_ and 510 μM K_2_SO_4_ for 5 days and then equal number of seedlings were transferred to 0 μM, 0.1 μM, 0.01 μM and 0.001 μM MeJA with pre-existing K_2_SO_4_ concentrations for 10 days. These seedlings were analyzed for root length, shoot length and lateral root length. After this analysis, the same seedlings were transferred to a hot air oven at 60°C for three days, and dry biomass was observed.

Arabidopsis (COL-0) seeds were surface-sterilized with 2% Sodium Hypochlorite solution for 1 min followed by washing with sterile water five times. After stratification at 4°, C in dark seeds were transferred to ½ strength Murashige and Skoog (1962) media and kept at 22°C and 16/18 hrs light/dark photoperiod. Equally germinated seedlings were transferred to ½ MS containing 9.4 mM KNO_3_ and 2.5 μM KI (K sufficient) and 2.5 μM KI (K deficient). The media was supplied with 0 μM, 0.1 μM, 0.01 μM and 0.001 μM MeJA dissolved in DMSO. Root growth was analyzed after five days of growth.

### Transcriptome Analysis

Uniformly germinated WT and *OsJAZ9* overexpressing line OE (OE10) seedlings were transferred to 4.08 μM and 510 μM concentrations of K_2_SO_4_ for 15 days along with other nutrients as per Yoshida Liquid media. After that, total RNA was isolated with Qiagen RNeasy Mini Kit from the whole seedling according to the manufacturer’s protocol. RNA integrity was analyzed using Bioanalyzer (2100 Agilent technologies), and samples with RIN value ≥7.5 proceeded for library preparation. Transcriptome sequencing was done using the Illumina Hiseq 2500/4000 platform with two independent biological replicates for each WT and *OsJAZ9* OE10. Adapter sequences and low-quality bases were trimmed using AdapterRemoval v2 (version 2.2.0). Further, rRNA sequences were removed by aligning reads with silva database. Finally, processed reads were aligned with the MSU7 genome (http://rice.plantbiology.msu.edu/pub/data/Eukaryotic_Projects/o_sativa/annotation_dbs/pseu domolecules/version_7.0/all.dir//) using the STAR program (Version 2.5.3a). Differentially expressed genes were identified as described (Bandyopadhyay et al., 2017).

### JA estimation

For Jasmonic Acid content estimation, equally germinated WT, *OsJAZ9* RNAi (RN12) and overexpression (OE10) seedlings were transferred to 4.08 pM and 510 pM concentrations of K_2_SO_4_ along with other nutrients as per Yoshida Liquid media for 15 days. JA concentrations in shoot samples were estimated in at least four replicates using Plant Jasmonic Acid (JA) ELISA Kit (MyBioSource, Inc.), following the manufacturer’s protocol.

### *R. solani* infection assay

*R. solani* AG1-IA isolates of BRS1 were used as a fungal pathogen. Fully grown *R. solani* sclerotia on Potato Dextrose Agar plates (PDA, Himedia, India) were used for pathogen inoculation on different rice plants. Sclerotia collected from PDA plates were incubated in 10 ml of Potato Dextrose Broth (PDB) for six h at 28°C to induce sclerotia germination. Tillers of 45 days old WT, *OsJAZ9* OE and RNAi transgenic rice plants, grown in soil (28°C temperature, 80% relative humidity and 16/8 photoperiod), were inoculated with fungal sclerotia. Samples of infected (*R. solani*) plants were collected at 3 and five days postinoculation (dpi) for further study. Relative vertical sheath colonization (RVSC) was calculated using the method described earlier (Ghosh et al. 2014). For cell death assays infected sheath samples were incubated with 0.02% Trypan blue reagent for 16 h at room temperature with continuous shaking. After that, samples were stained using distaining solution (3:1 ratio of glacial acetic acid and ethanol) followed by washing with distilled water. Samples were observed under a Nikon stereo-zoom microscope for blue color staining (Nikon AZ100).

## RESULTS

### K deficiency promotes higher expression of JA biosynthesis and signaling in rice

Recent transcriptome data suggest upregulation of a large number of JA associated genes under K deficiency (Ma et al., 2012; Takehisa et al., 2013; Shankar et al., 2013). We also noticed that root growth inhibition is a typical growth response to both JA treatment and K deficiency. This invoked us to look further into overlapping transcriptional responses of JA signaling modules under both JA treatment and K deficiency conditions. Further, we have previously reported that JA signaling repressor *OsJAZ9* is influenced by K deficiency (Singh et al., 2015) and is localized in the nucleus (Fig S1). To further explore the role of JA signaling under K deficiency, we analyzed temporal expression dynamics of various JA-associated genes including *OsCOI1b* (JA co-receptor), *OsAOS1*, *OsAOS2*, *OsJAR1* (JA biosynthesis) and *OsMYC2* (JA signaling) under K deficiency in rice. Interestingly, expression of *OsJAZ9* was found downregulated up to 6 h of the deficiency, and after that, it stabilized to normal. Induced expression of *OsCOI1b* and *OsMYC2* indicated enhanced JA signaling while increased expression of *OsAOS1*, *OsAOS2* and *OsJAR1* suggested enhanced JA signaling and biosynthesis under K deficiency (Fig 1). These results imply a higher intrinsic level of JA and signaling during the initial phase of K deficiency in rice.

**Figure 1.**
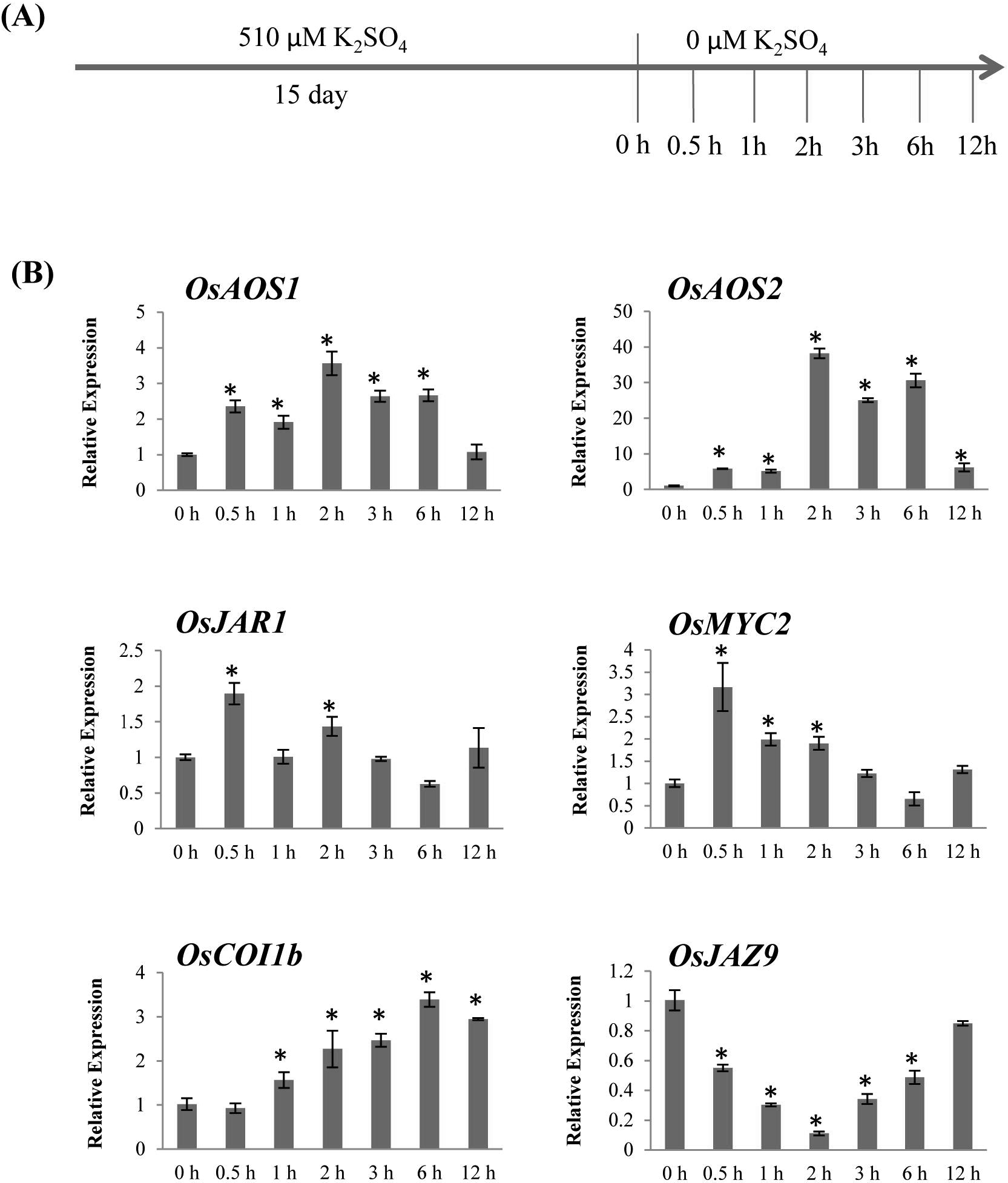
Potassium deficiency induces JA signaling in rice. **(A)** Schematic representation of experimental setup used for expression analysis of JA biosynthetic and signaling genes. Plants were raised under normal conditions (510 μM K_2_SO_4_) for 15 days before being subjected to different durations (0.5 to 12 h) of K deficiency treatment (0 μM K_2_SO_4_). **(B)** Short-term K deficiency responses of JA biosynthetic and signaling genes. Relative expression levels at different time points of K deficiency were analysed with respect to 0 h. Error bar indicates standard error derived from three independent replicates. Asterisk indicates significant changes at different time points compared to 0 h treatment (*p* ≤ 0.05, Student’s *t*-test).

We noticed previously (Singh et al., 2015) that among all rice *JAZs* genes, *OsJAZ9* was highly downregulated gene after 15 days of K deficiency. Also, *OsJAZ9* was explicitly most responsive towards K deficiency among N, P, K, Zn and Fe deficiency stresses. Therefore, we hypothesized that K deficiency needs increased JA signaling, and OsJAZ9 plays an important role.

To investigate it further, we generated *OsJAZ9* overexpression/RNAi and OsJAZ9ΔC translational reporter lines fused with GUS. *OsJAZ9:GUS* and *OsJAZ9ΔC:GUS*, (Full-length *OsJAZ9* and C-terminal deleted *OsJAZ9* fused with GUS, respectively) were cloned under 35S promoter (Fig 2A; S2; S3). Me-JA treatment promoted the degradation of GUS signal in OsJAZ9:GUS reporter roots; however, GUS signal was intact in the roots of OsJAZ9AC:GUS reporter line (Fig 2B) implying that OsJAZ9 is degraded in the presence of JA through Jas domain. Further, Me-JA supplemented with MG-132 (proteasome inhibitor) was unable to degrade OsJAZ9:GUS signal which confirms that JA degrades OsJAZ9 via proteasomal degradation pathway and this degradation is mediated by C-terminal Jas motif (Fig 2B). Therefore, using these two lines, we investigated JA response during K deficiency taking OsJAZ9:GUS stability as a proxy for JA action. Interestingly, the initial exposure of K deficiency (up to 3h) showed a rapid decrease in OsJAZ9:GUS signals. However, after 6h of K deficiency, OsJAZ9:GUS signal was stabilized indicating early induction of JA signaling under K deficiency. On the other hand, OsJAZ9ΔC:GUS line showed consistent GUS levels throughout the K deficiency (Fig 2C). This indicates a transient increase in intrinsic JA levels in roots during the initial phase of K deficiency. All these results confirmed that JA signaling/levels are indeed enhanced under K deficiency response in rice, and *OsJAZ9* is a critical component of this response.

**Figure 2.**
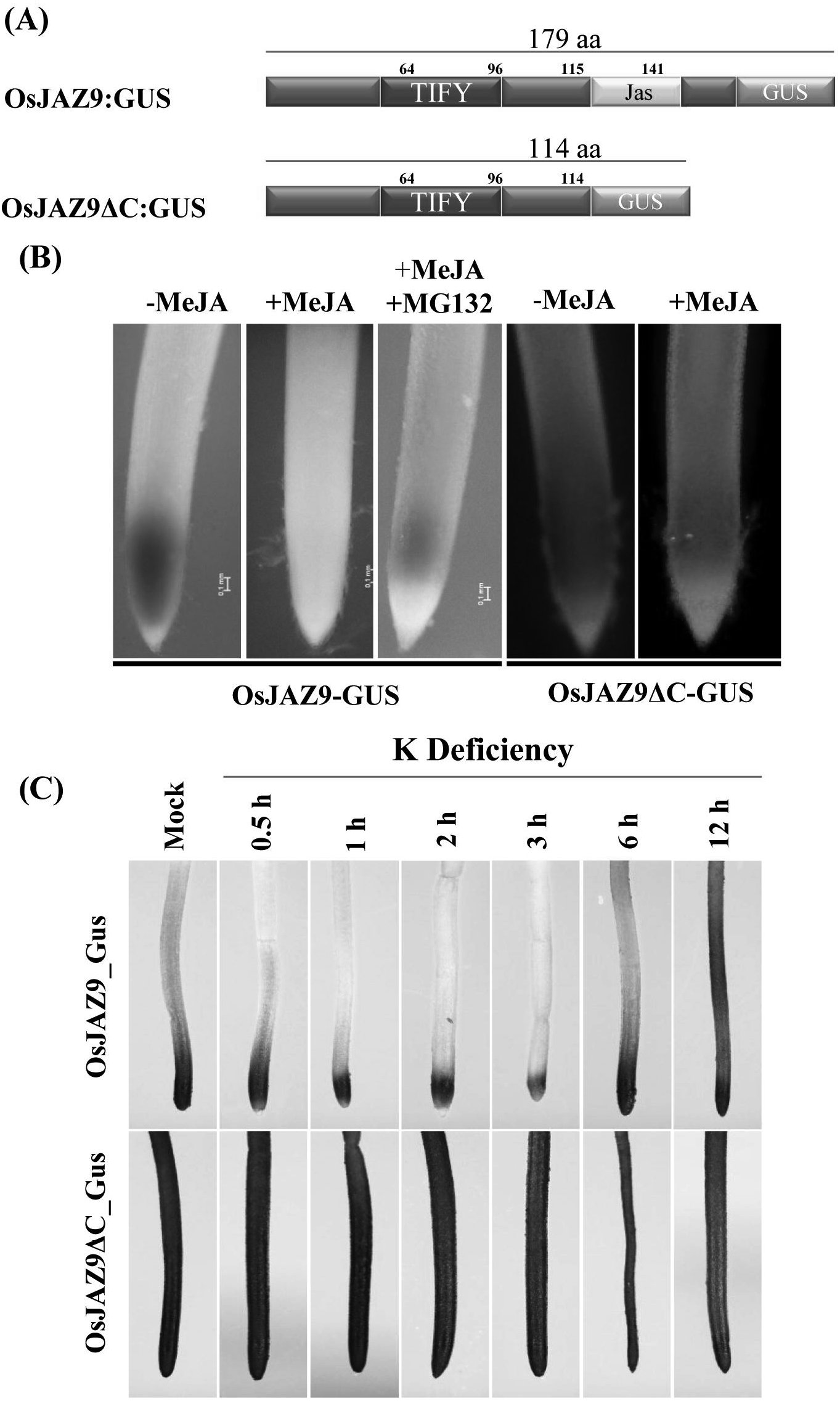
Potassium deficiency causes rapid increase in JA levels in rice. (**A**) Schematic representation *of OsJAZ9:GUS* and *OsJAZ9*-Δ*C:GUS* (Jas domain deleted) translational reporter constructs. **(B)** JA dependent proteasomal degradation of OsJAZ9 through Jas domain. Root tips of plants overexpressing *OsJAZ9:GUS* and *OsJAZ9*Δ*C:GUS* were analysed for GUS activity after 3h of treatment with 100 pM Me-JA alone or combined with 100 μM proteasome inhibitor, MG 132. **(C)** 15-days-old rice seedlings expressing *OsJAZ9:GUS* and *OsJAZ9ΔC:GUS* were transferred to K deficient media for indicated time points (0.5 to 12 h). Stability of OsJAZ9:GUS and OsJAZ9AC:GUS proteins in response to K deficiency were analyzed by histochemical GUS staining of root sections. Experiment was performed in three independent replicates.

### Overexpression of *OsJAZ9* enhances K deficiency tolerance in rice

The observed functional link between *OsJAZ9* and K deficiency responses made us analyze the growth behavior of *OsJAZ9* transgenic lines under K deficiency. Therefore, we generated homozygous lines of *OsJAZ9* overexpression (OE1, 3 and 15) and RNAi (RN12 and 14) lines. The overexpression lines had up to 120 fold upregulation while RNAi lines showed around a 98% reduction in the *OsJAZ9* expression as compared to WT plants (Fig S3). To examine the impact of K deficiency on plant growth, we first screened wild-type rice seedlings on a series of K concentrations (ranging from 0 to 510 μM) and found that 4.08 μM K_2_SO_4_ was able to inhibit ~50 percent of root growth as compared to 510 μM K_2_SO_4_ (Fig S4). Therefore, we used this concentration of K_2_SO_4_ as K deficient media for further experiments. Phenotypic analysis revealed that overexpression of *OsJAZ9* helped plant gain higher shoot biomass under K deficiency while in normal conditions no significant difference was observed in the shoot biomass of OE, RNAi and WT plants (Fig S5). Interestingly, root biomass of *OsJAZ9* OE lines was highly reduced during K deficiency while RNAi lines were almost similar to WT roots (Fig S5). We also analyzed growth behavior of *OsJAZ9:GUS* OE lines (OE2 and OE10; Fig S6) under K deficiency. These *OsJAZ9* overexpressing lines also showed increased shoot biomass under K deficiency (Fig S6). With these results, we conclude that overexpression of *OsJAZ9* imparts K deficiency tolerance to rice plants.

To further analyse whether OE plants have higher K use efficiency or K uptake efficiency, we examined growth behaviour of WT, RNAi (RN/14), OE (OE/10) and *OsJAZ9ΔC* OE (J9ΔC) plants grown under K sufficient conditions for 10 days (T1) followed by K deficient conditions (0 μM K_2_SO_4_) for 15 days (T2) and recovery with 10 μM K_2_SO_4_ for 5 days (T3). *OsJAZ9 OE* line usually showed reduced root and shoot growth as compared to WT, but under K deficiency and recovery conditions, it showed better growth performance (Fig 3). This change becomes more apparent with J9ΔC OE line which usually does not have any difference in either shoot and root growth and remains non-significant during K deficient and sufficient conditions, but it showed significantly faster growth during recovery conditions (Fig 3). These results indicated that *OsJAZ9* overexpression plants are adapted for both better K use efficiency (*OsJAZ9* OE plants were able to produce more biomass during zero external K concentrations) and K uptake efficiency (*OsJAZ9* OE plants produced faster growth during recovery). Interestingly, we also observed higher Na^+^/K^+^ in OE plants then both WT and RNAi during K deficient conditions. This could be explained well as Na+ and K+ are the structural mimics of each other and Na^+^ is taken up when K^+^ is not available to the plants which can result into higher Na^+^/K^+^ ratio under extremely K deficient conditions (0 μM K) (Fig 3). This becomes clearer when the K starved plants were recovered with 10 μM K_2_SO_4_ and Na^+^/K^+^ goes down in OE lines, but it remains high in RNAi lines. This decrease in the ratio of Na^+^/K^+^ was more due to the efficient K uptake rather Na uptake by OE lines as indicated by the K+ content per plant.

**Figure 3.**
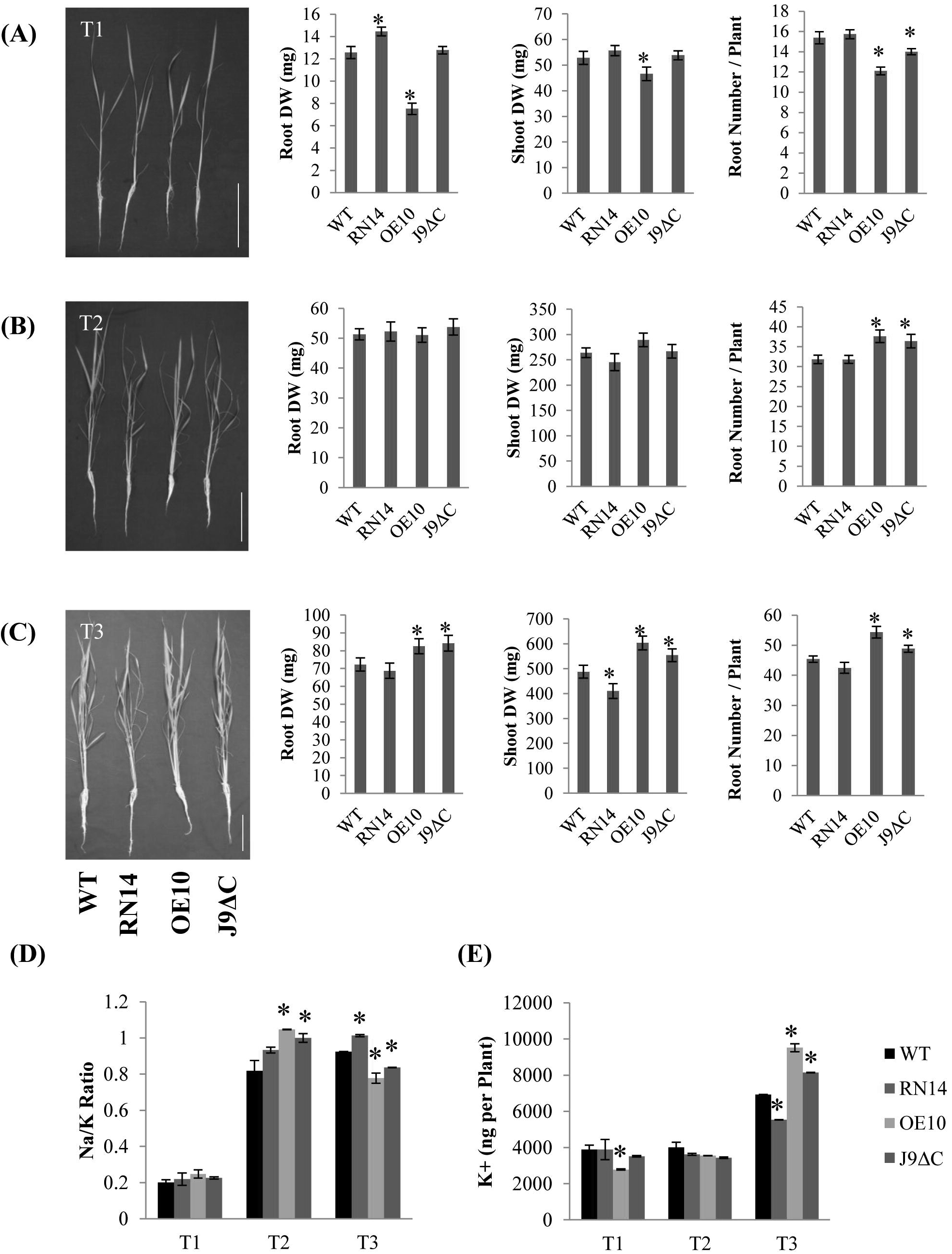
Enhanced K acquisition in K-starved *OsJAZ9* OE seedlings upon K resupply. **(A)** Morphological data (root dry weight, shoot dry weight and number of roots per plant) of 10-days-old normally grown seedlings (T1), **(B)** followed by 15-days of K starvation (T2) and **(C)** subsequent recovery with 10 μM K_2_SO_4_ for 5 days (T3). **(D)** Na^+^/K^+^ and **(E)** Total K content per plant of WT and *OsJAZ9* transgenics during T1, T2 and T3 treatments. For biomass measurements, bar represents average of 15 seedlings with standard error. K content experiment was performed with two replicates. Asterisks indicate significant changes in transgenics compared to WT at respective time points (*p* ≤ 0.05, Student’s *t*-test). Scale =10 cm.

### *OsJAZ9* OE modulates JA signaling and K acquisition in rice

To understand the basis of improved K uptake of *OsJAZ9* OE transgenic lines, we analyzed the expression pattern of known K transporter genes in WT seedlings. Most of the genes coding for K transporters (including *OsHAK2;1, OsHAK5* and *OsHAK1)* were induced on K deficiency. Further, JA signaling gene (OsMYC2, OsCOI1b) also followed the same patterns confirming the level of K deficiency experienced by the plants and activation of JA signaling (Fig S7A). Interestingly, we did not find *OsNHX1* induction under K deficiency while its transcripts were induced during salt stress (Wu et al., 2015). Thus its non-responsive nature towards K deficiency demarcates the thin boundary between salt and K deficiency stress. Next, we investigated how these genes behave in *OsJAZ9* transgenic lines. Interestingly, we found higher expression of *OsHAK2;1* and *OsHAK1* in *OsJAZ9* OE lines especially after 12 h of K recovery (Fig S7B).

### Overexpression of *OsJAZ9* affects JA signaling in rice

Overexpression of *JAZs* under constitutive promoter has resulted in JA increased insensitivity in Arabidopsis and rice previously (Thines et al., 2007; Yamada et al., 2012). Therefore, we analyzed root growth inhibition in *OsJAZ9* OE and RNAi lines on MeJA treatment. Consistent to previous studies, we also observed a partial JA insensitive phenotype in *OsJAZ9* OE lines (Fig 4). We observed ~57% root growth inhibition in RNAi line (RN12), 45% in WT, 27% and 12% change (inhibition) in OE lines, OE3 and OE10, respectively on MeJA application. Therefore, this confirms that *OsJAZ9* is capable of modulating JA signaling or response.

**Figure 4.**
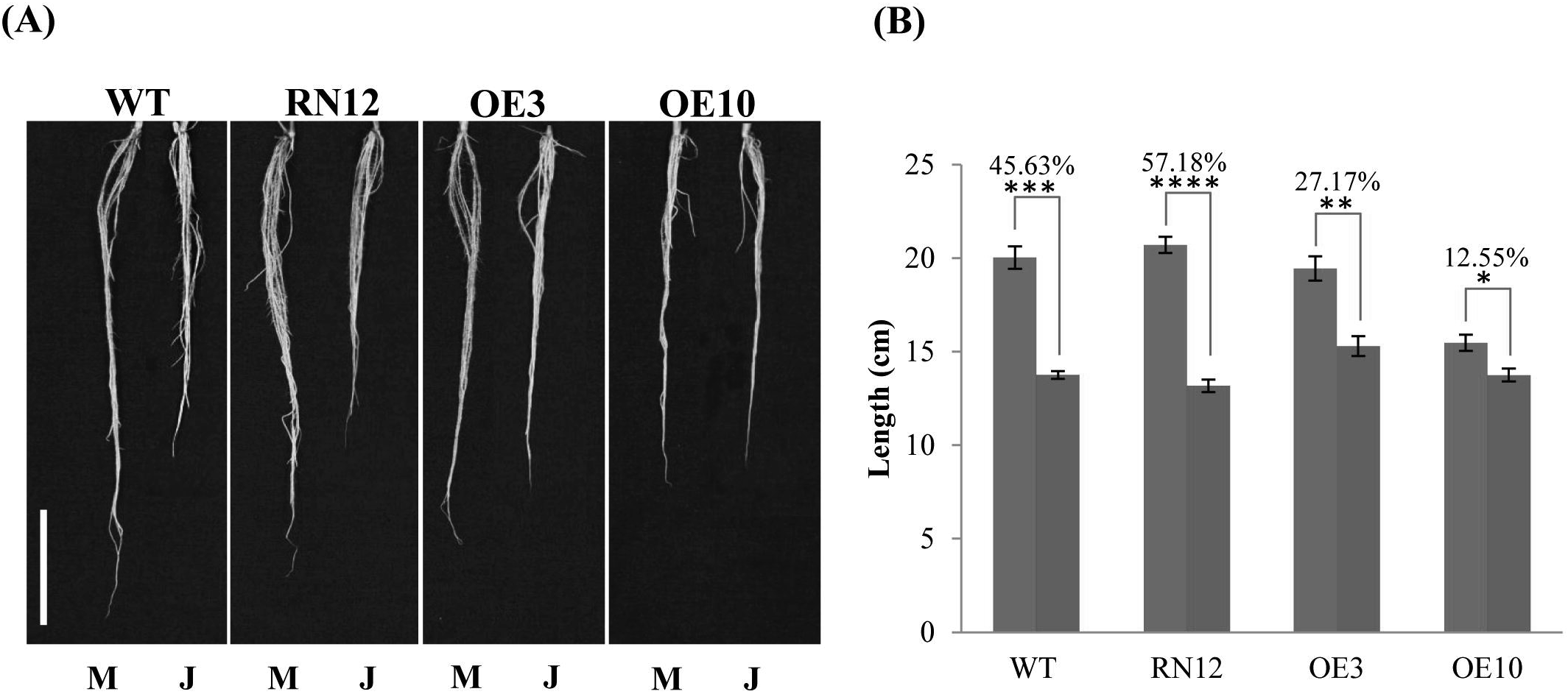
JA-induced root inhibition in WT and *OsJAZ9* transgenics. **(A)** Representative images of roots of 20-days old WT, RN12, OE3 and OE10 seedlings supplemented with DMSO (Mock; M) or 10 μM MeJA dissolved in DMSO (JA treated; J). Scale = 5 cm. **(B)** Average root lengths of WT, *OsJAZ9* RNAi and OE plants under DMSO (blue bars) and MeJA (Red bars) treatments. Each bar represents average of 9 seedlings with standard error. Values over the bars represent JA mediated percentage reduction in the root length. Significant differences between DMSO vs MeJA treatment were evaluated by Student’s *t*-test. Asterisks; *, **, *** and **** indicate *p* values, ≤ 10^−3^, 10^−4^, 10^−8^ and 10^−10^, respectively.

We next performed RNA-seq analysis of WT and *OsJAZ9* overexpression transgenic lines to uncover the effects on JA signaling and genes involved in K deficiency tolerance. WT and *OsJAZ9* OE (OE/10) line raised under normal and K deficient conditions for 15 days were used in this study. A total of 5173 and 5505 genes were differentially expressed (p<0.05, q<0.05) in WT and *OsJAZ9* OE line, respectively, during K deficiency. Out of these 2433 and 2670, genes were upregulated, and 2740 and 2835 genes were downregulated, respectively, in WT and *OsJAZ9* OE line (Fig 5). This analysis shows that *OsJAZ9* OE lines are more responsive towards K deficiency as compared with WT plants.

**Figure 5.**
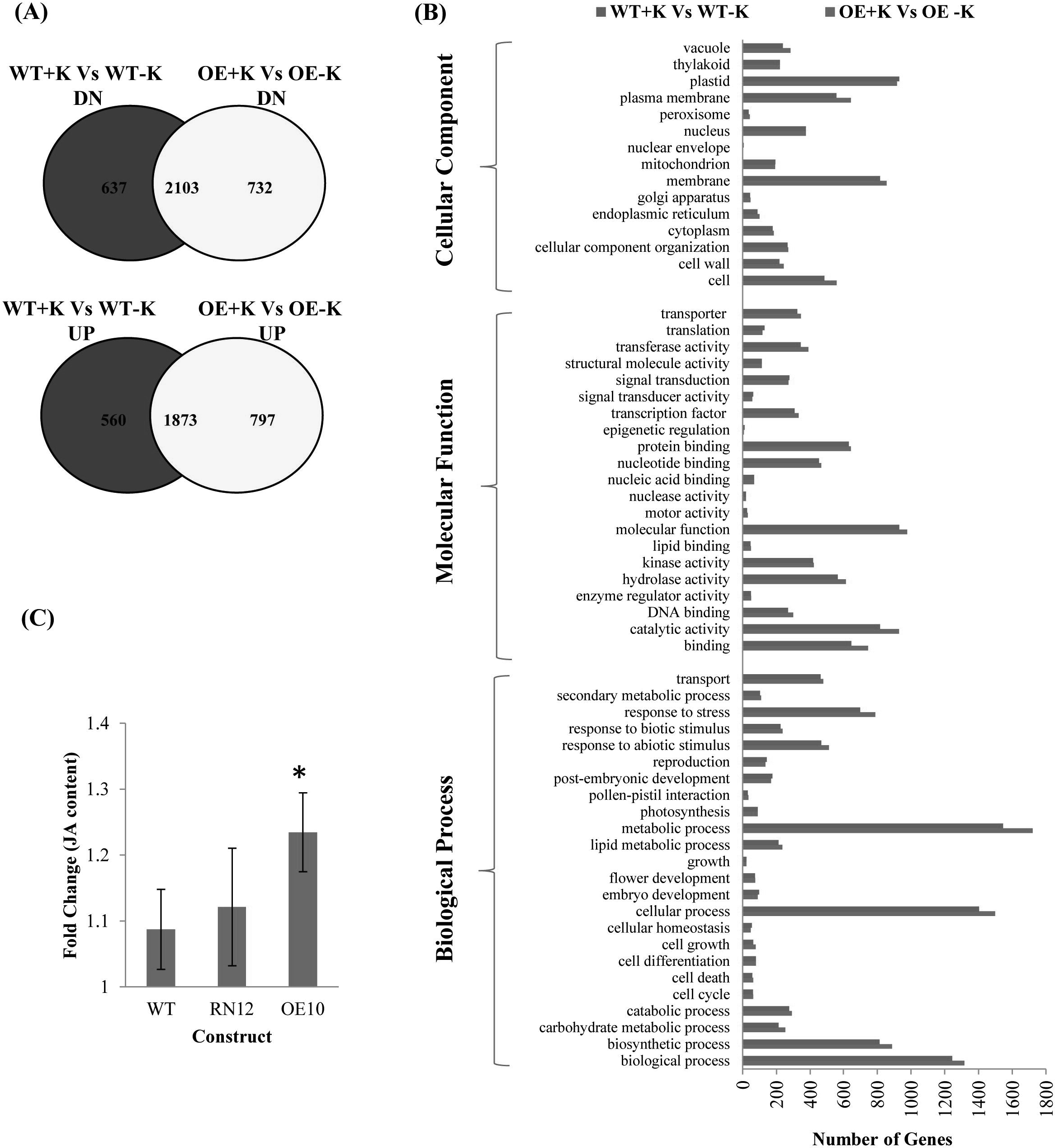
Overexpression of *OsJAZ9* enhances K deficiency response and JA content in rice. **(A)** Venn diagrams showing differentially regulated genes on K deficiency in WT and OsJAZ9 OE line. **(B)** Overview of transcriptome data showing category-wise distribution of differentially genes in WT and *OsJAZ9* OE lines during K deficiency. **(C)** Fold change in JA content in WT, OsJAZ9 RNAi (RN12) and OE (OE/10) lines during 15 days of K deficiency. Bars represent the average change in JA content among at least four replicates. Error bar represents SE of all the replicates. (*p* ≤ 0.05, Student’s *t*-test).

We further analyzed the expression of all known JA biosynthesis and signaling genes in our transcriptome data. We observed that out of 17 JA- biosynthesis related genes detected in the transcriptome, seven were having higher abundance in OE plants as compared to WT during K deficiency considering their respective controls while four were downregulated (Table S2). Upregulated genes included *OsAOS2, AOS3, OsLOX5, OsLOX7, OsLOX8, OsACX2, OsOPR5,* and *OsJAO1.* We also found higher expression of key JA marker genes like *OsLEA3* and *OsVSP2* in *OsJAZ9* OE plants than WT plants due to K deficiency (Table S2). This higher abundance of JA biosynthesis, signaling and marker genes confirms the higher JA signaling in *OsJAZ9* OE plants during K deficiency as compared to the WT plants. Further Analysis revealed that out of 13 known K transporters, 7 were having higher abundance in OE line as compared to WT in response to K deficiency, while 6 transporters were expressed less. Among the lesser expressed ones, *OsHKT27* codes for a K efflux transporter, OsHKT2;2 and OsHKT1;5 are more efficient in Na transport than K (Kadar et al., 2006) while OsK1.1 (OsAKT1) functions at the millimolar range of external K (Ahmad et al., 2016). We found *OsHAK10, OsHAK24, OsK3.1, OsK5.2, OsHKT1;1, OsHKT1;4, OsSOS1* and *OsGORK* genes having higher abundance in OE line as compared to WT in K deficient conditions and thus supports the case for higher K uptake in *OsJAZ9* overexpressing plants (Table S3).

### *OsJAZ9* modulates root system architecture

As overexpression of *OsJAZ9* results in higher K uptake, we analyzed root phenes that can contribute to K uptake in addition to the transporter activity. Interestingly, *OsJAZ9* OE plants showed shorter seminal roots while RNAi (RN14) plants have longer seminal roots both in K deficient and sufficient conditions. However, *OsJAZ9* OE lines had higher lateral root length (Fig 6). It seems while seminal root length is inhibited in OE lines, increase in laterals length might have increased root surface area for better K acquisition. Further, this phenotype seems to be *OsJAZ9* dependent as we observed a similar phenotype under both K deficient and sufficient conditions.

**Figure 6.**
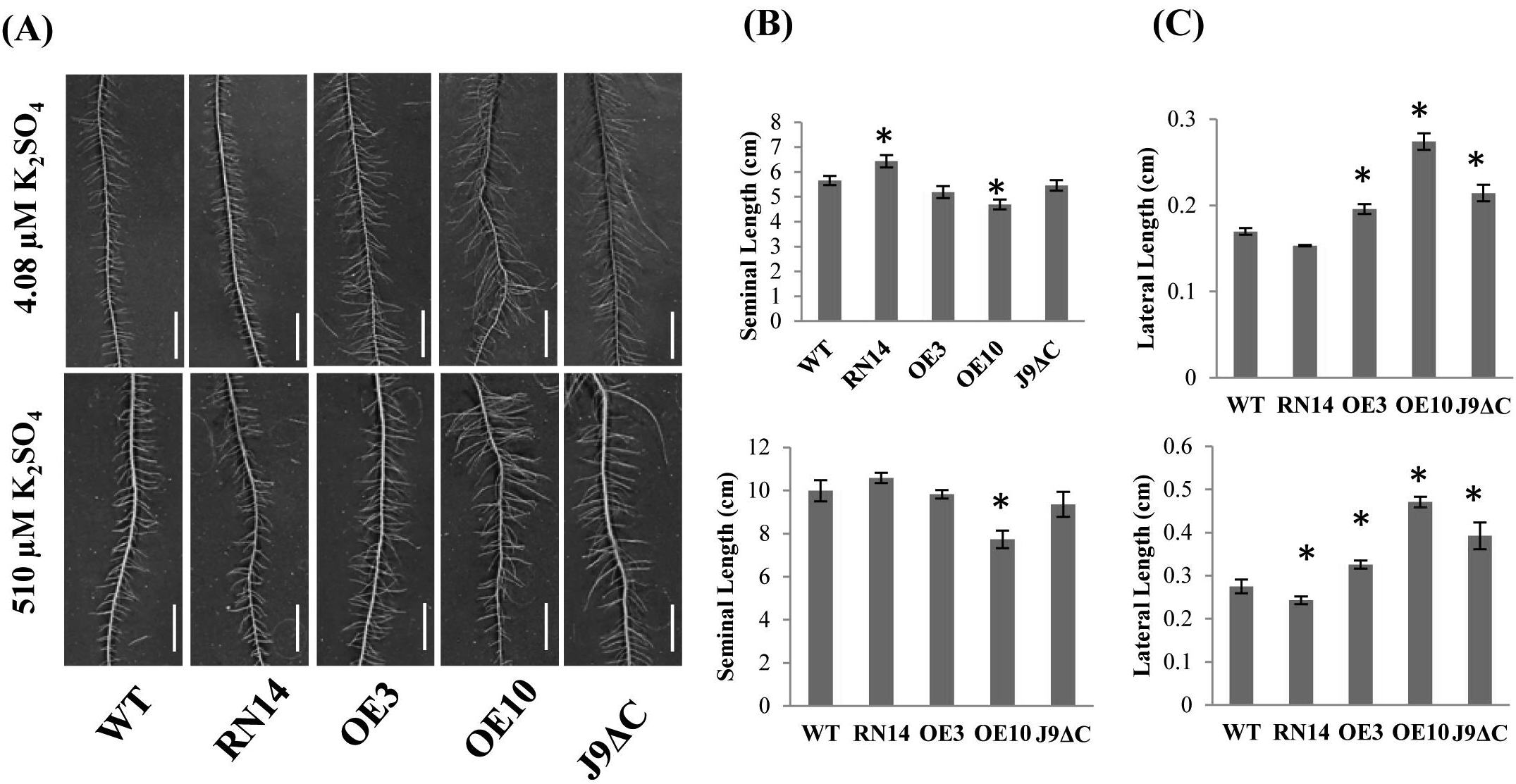
*OsJAZ9* overexpression influences root system architecture. **(A)** Representative root images of 12-days-old WT and *OsJAZ9* transgenics showing lateral root lengths during K deficient (upper panel) and K sufficient conditions (lower panel). **(B)** Average seminal root length and **(C)** Average lateral root length of WT and *OsJAZ9* transgenics during K deficient and K sufficient conditions. Four independent plants were used for the analysis. Scale bar represents 1 cm while error bar represents SE among the replicates. (*p* ≤ 0.05, Student’s *t*-test).

Previously, JA treatment at lower concentrations was found to enhance the lateral length in Arabidopsis and rice, though at higher levels it inhibits lateral length (Raya-Gonzalez et al., 2012; Hsu and Kao, 2011). We speculated that this shortening of root length along with enhanced lateral length is caused by higher levels of intrinsic JA or due to enhanced JA signaling in *OsJAZ9* OE lines. To validate this, we analyzed the effect of exogenous MeJA application on lateral root length. Interestingly, we observed increased lateral root length at lower concentrations of MeJA (0.001 and 0.01 μM) while the higher level of MeJA (1 μM) inhibited the lateral root length (Fig S8A and S8B). In agreement with Raya-Gonzalez et al. (2012) and Hsu and Kao (2011), we also observed increased lateral root numbers at lower concentrations of MeJA (Fig S8C). Again, longer laterals roots were overrepresented in with lower MeJA treatment as compared to no MeJA or higher MeJA concentrations (Fig S8D). Hence, the phenotype observed in *OsJAZ9* OE lines can also be achieved by treating WT plants with MeJA. These observations validate the hypothesis that *OsJAZ9* OE plants are experiencing higher JA signaling which is also reflected in transcriptome data.

### Overexpression of *OsJAZ9* enhances in-vivo JA levels under K deficiency

All these phenotypes hinted on the higher intrinsic JA levels or the increased JA signaling in *OsJAZ9* OE lines as compared to the WT. To check whether there was indeed higher JA content in *OsJAZ9* OE plants or these plants are only experiencing higher JA signaling we analyzed the change in the JA levels in WT, RNAi (RN12) and overexpression (OE/10) plants during K deficiency as compared to their respective controls. As suspected, WT and RNAi lines did not have any significant difference in the JA accumulation while OE line had a significant higher JA accumulation on K deficiency (Fig 5). This higher JA production in OE lines is well correlated with the higher induction of JA biosynthesis genes in OE lines under K deficiency.

As previously reported by Li et al. (2017) and Troufflard et al. (2010), we also observed induction in JA content due to K deficiency though the increase is meager which may be due to the prolonged period of the deficiency, i.e. 15 days or the milder K deficiency conditions as compared to the conditions used in previous reports. Here we conclude that overexpression of *OsJAZ9* results in enhanced natural JA levels which are also reflected in the transcriptome and morphological data.

### Low doses of exogenous MeJA helps plant tolerate K deficiency

To test whether exogenous JA application can help plant tolerate K deficiency, we analyzed growth behavior of rice seedlings under different concentrations of MeJA (0 μM, 0.1 μM, 0.01 μM, and 0.001 μM) supplied with K sufficient (510 μM K_2_SO_4_) and deficient media (4 μM K_2_SO_4_ and 8 μM K_2_SO_4_). We found an increase in root length and plant biomass by low concentrations of MeJA (<1 μM) during K deficient conditions (4 μM and 8 μM K_2_SO_4_) (Fig 7A and 7B). MeJA treatment at higher concentration (>1 μM) always inhibited all growth parameters studied during both low and high K conditions. At normal K concentrations, we observed either nonsignificant change or reduction in root length, shoot length and biomass by MeJA treatment (Fig 7C). We also noted that at severe K deficiency, a higher concentration of MeJA is required to overcome the deficiency symptoms. We also reconfirmed these results with Arabidopsis with different levels of MeJA supplied on K deficient media and found MeJA mediated growth inhibition in Arabidopsis under normal K concentration (Fig S9). Interestingly, Arabidopsis also showed better growth under K deficiency when supplied with low levels of MeJA; the further increase in MeJA concentration (0.1 μM) results in growth inhibition (Fig S9). These results confirm that low dosages of MeJA can help plant tolerate K deficiency.

**Figure 7.**
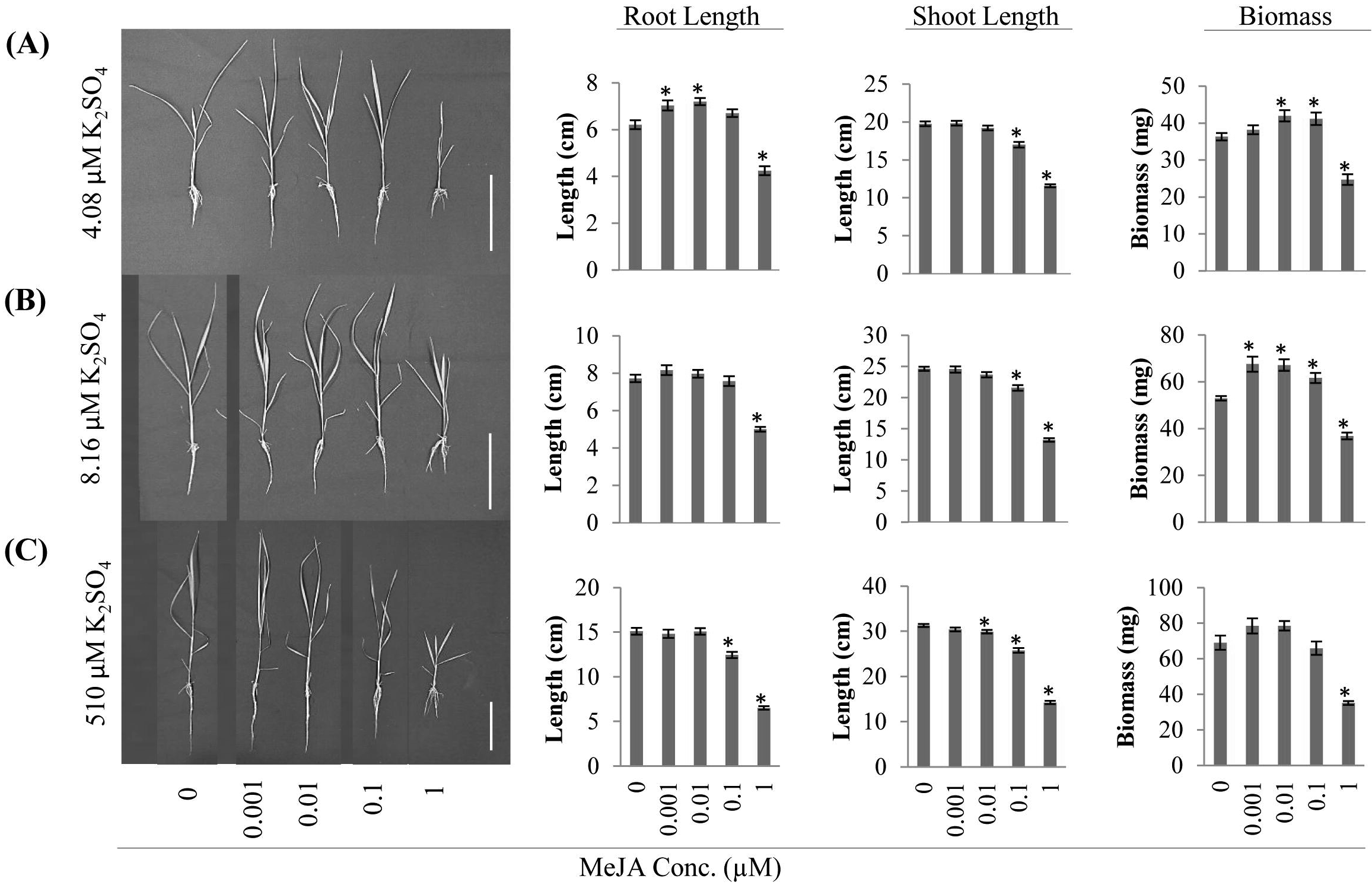
Exogenous application of MeJA treatment can help plant tolerate K deficiency. Plant growth after 15 days at (**A)** 4.08 pM K_2_SO_4_, (**B)** 8.16 μM K_2_SO_4_, and (**C**) 510 μM K_2_SO_4_ with different concentrations (0, 0.001, 0.01, 0.1 and 1.0 μM) of MeJA. Graphs in the respective rows show the average root length, Shoot length and biomass of 15 plants. Asterisks indicate significant changes in transgenics compared to WT at respective time points (p ≤ 0.05, Student’s t-test). Scale = 10 cm.

### *OsJAZ9* overexpression enhances sheath blight resistance in rice

Since manipulation of JA biosynthesis and signaling can have profound consequences on biotic stress tolerance of plants, we analyzed the influence of *OsJAZ9* on resistance to prime rice pathogen, *Rhizoctonia solani* which causes sheath blight disease. A previous report has shown that *OsWRKY30,* which is a JA responsive gene, can impart tolerance towards necrotrophic fungus *Rhizoctonia solani,* by increasing endogenous JA levels in rice plants (Peng et al., 2012). We have established that *OsJAZ9* overexpression enhances JA biosynthesis and JA signaling during low K stress. Therefore, we analyzed whether the endogenous accumulation of JA and enhanced JA signaling in *OsJAZ9 OE* plants results in *R. solani* resistance in rice. Interestingly, we observed lesion development and relative vertical sheath colonization (RVSC) index to be significantly less in *OsJAZ9* OE (OE3, OE10, J9ΔC) plants as compared to RNAi (RN12 and RN14) and WT plants (Fig 8A/B, Fig S10A/B). Being a necrotrophic pathogen, *R. solani* is known to impart cell death response in the infected tissues and therefore, we monitored the extent of host cell death in different transgenic lines by trypan blue staining. In case of infected WT and *OsJAZ9* RNAi lines the degree of visible blue color, which reflects host cell death, was found significantly higher as compared to the *OsJAZ9* OE (OE3, OE10, and J9ΔC) plants (Fig 8C, Fig S10C). Overall this data indicate that *OsJAZ9* OE lines were tolerant against *R. solani* infection while WT and RNAi lines were susceptible.

**Figure 8.**
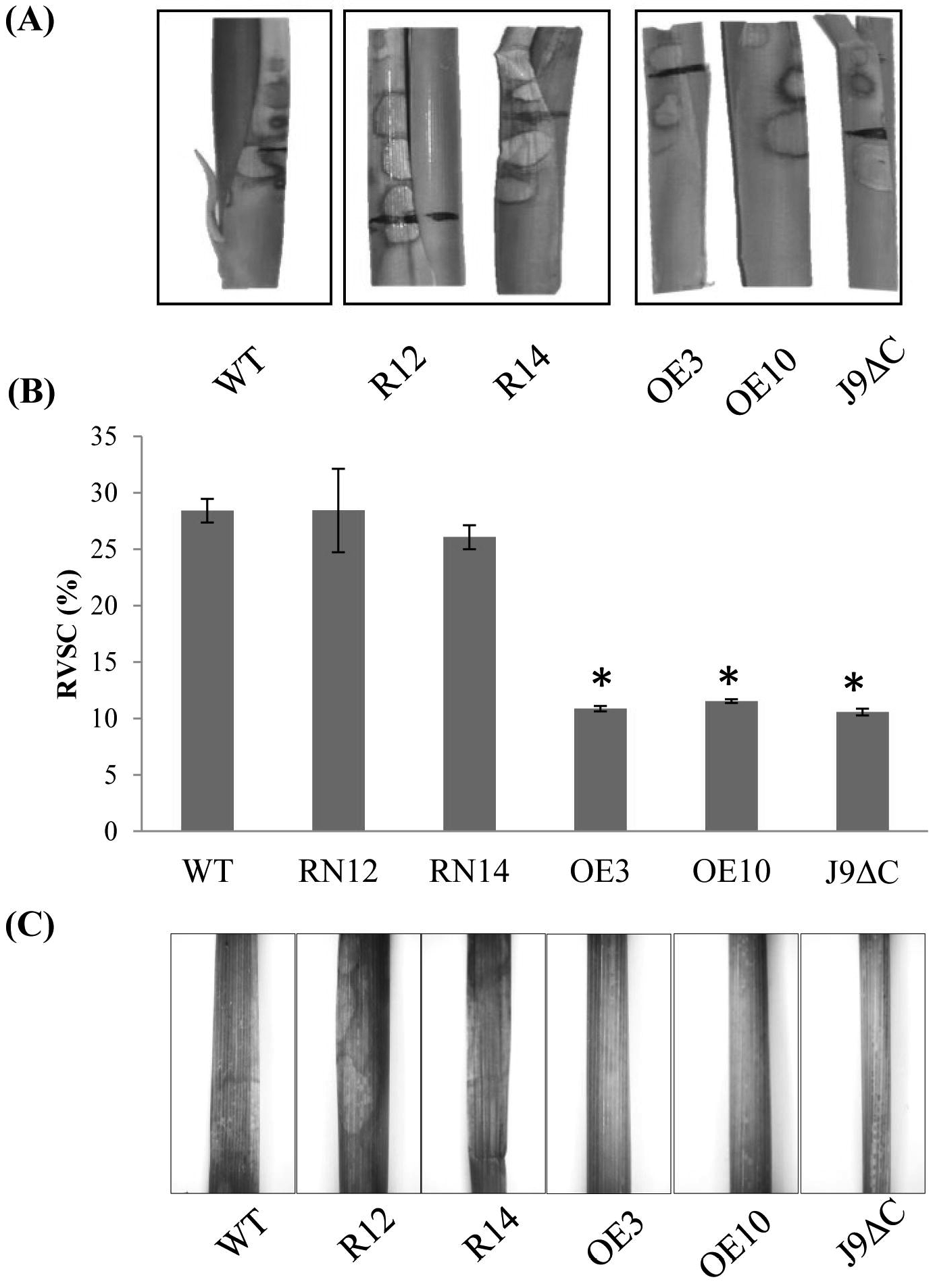
*OsJAZ9* overexpression imparts sheath blight tolerance in rice. **(A)** Representative images showing the extent of lesion development in WT, RN12, RN14, OE3, OE10 and J9ΔC lines at 5 days post-infection (dpi) by *Rhizoctonia solani.* **(B)** RVSC disease index indicating the severity of the infection after 5 days post infection. Tillers of 45-days-old plants were subjected to *R. solani* infection and disease symptoms were analysed at 5 dpi. Ten tillers (n=10) of each line were considered for infection. Error bar represents SE of all the replicates. (p ≤ 0.05, Student’s t-test). **(C)** Representative images of trypan blue stained leaf sheaths of infected tillers showing the extent of cell death caused by *R. solani* infection in WT and *OsJAZ9* transgenics.

## DISCUSSION

Potassium being a macronutrient is one of the most important minerals for plant growth. Despite its high concentration in soil, it remains mostly unavailable for root uptake and utilization. Therefore, its deficiency in soil remains one of the major constraints for crop productivity. As K^+^ being a structural mimic of Na^+^, its deficiency results in high Na^+^ uptake by plant and cause salt stresses. Therefore, K availability in the soil also controls the level of salt stress experienced by the plant. Further, K deficient plants are more susceptive to various biotic and abiotic stresses (Egilla et al., 2001; Hardter, 2002; Kant and Kafkafi, 2002; Kaya et al., 2006; Sarwar, 2012; Wang et al., 2013). Unlike Phosphorus deficiency response where enzymes like phosphatase can elevate the P availability in soil (Mehra et al., 2017), there is no enzyme known in rice to hydrolyze locked-up K in the soil. The strategies targeting K transporter and root architecture are more suited for enhancing K uptake efficiency in rice. While JAs are well-known for controlling plant defense against necrotrophic fungus and insects (Peng et al., 2012; Wasternack and Hause, 2013), emerging evidence has established their role in diverse plant processes like flower development, reproduction, response to abiotic and biotic stresses (Cai et al., 2014; Kazan, 2015). JAs are also known for inhibiting root growth for many years, and recently intricate details of this response were revealed (Wasternack and Hause, 2013; Wasternack and Song, 2017). Root growth patterns during K deficiency mimic the root phenotype observed on JA treatment in various plants (Staswick et al., 1992; Gruber et al., 2013; Cai et al., 2014). Recently, Troufflard et al. (2010) and Li et al. (2017) have found enhanced JA levels during K deficiency in Arabidopsis, rice, and wheat. JAZ repressors are pivotal to JA signaling, and their manipulation leads to many altered plant processes (Wasternack and Song, 2017). We have reported that *OsJAZ* genes coding for JA signaling repressors are transcriptionally influenced under K deficiency (Singh et al., 2015). However, it has not been demonstrated that how JAZs are involved in regulation of K deficiency response. Here, using comprehensive transgenics approach, we showed that *OsJAZ9* is a crucial component of JA signaling for K deficiency response (Fig 2).

Interestingly, *OsJAZ9* OE lines performed better in terms of root and shoot growth under K deficiency, while RNAi lines performed poorly. Moreover; low Na^+^/K^+^ found in OE lines was mainly because of increased K^+^ uptake as Na^+^ contents were less affected. These results suggested that *OsJAZ9* has improved K^+^ absorption over Na^+^ uptake during low K^+^ availability. The higher K+ absorption of *OsJAZ9* OE lines was well supported by the higher transcript abundance of K transporters, *OsHAK1* and *OsHAK2;1.* A higher magnitude of transcriptome changes in *OsJAZ9* OE lines further also showed greater responsiveness towards K deficiency than WT. Further, the enhanced expression of JA biosynthesis and signaling genes like *OsAOS2/3, OsLOX5/7/8, OsJAZO1* and *OsLEA3, OsVSP2*, respectively, confirms the higher JA signaling in *OsJAZ9* OE plants during K deficiency. The enhanced JA signaling on overexpression of a repressor is surprising; however, similar observations on activated JA signaling were made on *JAZ7* overexpression in Arabidopsis (Thatcher et al., 2016). In addition to K transporters, longer laterals might have also contributed to improved K uptake in *OsJAZ9* OE plants. Previously, exogenous MeJA has resulted in the enhanced lateral root length and density in Arabidopsis (Raya-González et al., 2012; Hsu and Kao, 2011). Our observations of lateral root induction by lower concentrations of MeJA confirm growth-promoting nature of JA at a low level and complement the longer lateral phenotype of *OsJAZ9* OE lines. MeJA mediated K deficiency tolerance in rice and Arabidopsis observed in our data further support this hypothesis. However, it may hold true only at lower levels of JA as higher exogenous JA is growth inhibitory irrespective of the stress. These increased JA signaling or intrinsic levels of JA can also have other stress-related advantages for plants. Previously, Peng et al. (2012) argued that higher endogenous JA levels result into tolerance towards necrotrophic fungus *R. solani.* Tolerance against *R. solani* infection by *OsJAZ9* OE in our experiment confirmed this argument and supported increased JA signaling or the higher levels of JA on *OsJAZ9* overexpression.

At morphological levels, JA treatment is known to inhibit the primary root growth in plants (Staswick et al., 1992; Cai et al., 2014). Consistent with this phenomenon, we found roots of *OsJAZ9* OE plants shorter than WT; mimicking the JA overproduced or enhanced JA signaling response. This phenotype is very appealing as most of the other *JAZ* repressors overexpressed transgenics in Arabidopsis (*JAZ1, JAZ2, JAZ3, JAZ5, JAZ7,* and *JAZ9*) and rice (*JAZ1* and *JAZ9*) reported earlier, either promoted shoot growth or showed no change (Chini et al., 2007; Hakata et al., 2017; Yang et al., 2012). Like other JAZs, root growth inhibition assay revealed reduced JA sensitive behavior of *OsJAZ9* OE lines while RNAi lines were JA hypersensitive. The induction of JA biosynthetic genes and the phenotypes mimicking the MeJA treatment made us analyze the JA content in our transgenic lines. The slightly enhanced JA accumulation in *OsJAZ9* transgenic lines explains higher JA signaling in OE lines during K deficiency. Mostly, overexpression of Arabidopsis JAZs inhibits JA induced growth suppression while the loss of function mutants of *AtJAZs* exhibited hypersensitivity to JA (reviewed in Wasternack and Song, 2017). In agreement to Wu et al. (2015), we also found JA hypersensitive response in *OsJAZ9* knockdown lines. However, elevated JA levels and JA signaling in *OsJAZ9* OE lines are very intriguing. We suspect here a negative feedback regulation of JA homeostasis wherein constitutive overexpression of a JAZ repressor hyperactivated JA signaling (Chini et al., 2007; Thatcher et al., 2016). JAZs activities are tightly controlled at tissue levels. By constitutively overexpressing a JAZ repressor we may have disturbed that finely tuned JA homeostasis resulting in enhanced JA levels and signaling in *OsJAZ9* overexpression plants as also reported earlier in Arabidopsis (Thatcher et al., 2016). Noticeably, JA levels were also enhanced in knockdown lines of *AJAZ1* while remaining unchanged in the knockdown lines of *NaZAZh* (Li et al., 2017; Oh et al., 2012). Further research is needed to pinpoint the chief regulator of JA negative feedback regulation on JAZ overexpression. A closer look in the transcriptome data revealed the suppression of all JAZ repressors in under both normal and low K conditions in *OsJAZ9* overexpression plants (Table S4). This would make OsMYC2 free from the clutch of JAZ repressors which leads to enhanced JA signaling which in-turn shape overall phenotype and its effect in root growth. The suppression of all other JAZs can underscore the only suppression mediated by OsJAZ9 overexpression. However, how low K level at cellular resolution in cellular milieu invokes JA biosynthesis and signaling will pave the way for a broader understanding of low K perception and its close association with JA machinery. Nevertheless, overexpression of *OsJAZ9* was reported to impart salt tolerance via modulating K transporters in rice (Wu et al., 2015). Our work establishes its role in K deficiency tolerance and revealed novel aspects of JA mediated response via JAZ repressors.

## ACKNOWLEDGMENTS

APS, PM, and BKP acknowledge the research fellowship by UGC, CSIR, and DBT, respectively. We thank Dr. Vijayata Singh, CSSRI, Karnal for help in K and Na estimation. JG acknowledges a grant from INSA-young scientist project.

## AUTHORS CONTRIBUTIONS

APS, BP, and PM designed and conducted experiments. APS, RK, and GJ conducted *R. solani* infection experiment. JG designed project supervised experiments and analyzed data. APS, BP, PM, and JG wrote the manuscript.

## Supplementary Data

**Figure S1.** Nuclear localization of YFP-OsJAZ9.

**Figure S2.** Protein sequence alignment of OsJAZ9 and OsJAZ9ΔC.

**Figure S3.** Raising and screening of *OsJAZ9* rice transgenics.

**Figure S4.** Effect of K deficiency on rice growth.

**Figure S5.** *OsJAZ9* is involved in K deficiency tolerance in rice.

**Figure S6.** Effect of *OsJAZ9* overexpression on growth parameters of rice during K sufficient and deficient conditions.

**Figure S7.** Overexpression of *OsJAZ9* enhances expression of K transporters.

**Figure S8.** Exogenous application of MeJA modulates root system architecture.

**Figure S9.** Growth promoting effects of lower concentration of JA under K deficiency.

**Figure S10.** *OsJAZ9* overexpression imparts sheath blight tolerance in rice.

**Supplementary Table S1.** List of primers used for qRT-PCR and gene cloning.

**Supplementary Table S2.** Effect of *OsJAZ9* expression on expression of JA associated genes under K deficiency.

**Supplementary Table S3.** Effect of *OsJAZ9* expression on K transporter expression under K deficiency.

**Supplementary Table S4.** Effect of *OsJAZ9* expression on expression of *OsJAZ* genes.

## References

Ahmad I, Mian A, Maathuis FJM. 2016. Overexpression of the rice AKT1 potassium channel affects potassium nutrition and rice drought tolerance. Journal of Experimental Botany 67: 2689–2698.

Armengaud P, Breitling R, Amtmann, A. 2004. The potassium-dependent transcriptome of Arabidopsis reveals a prominent role of jasmonic acid in nutrient signaling. Plant Physiology 136: 2556–2576.

Bandyopadhyay T, Mehra P, Hairat S, Giri J. 2017. Morpho-physiological and transcriptome profiling reveal novel zinc deficiency-responsive genes in rice. Functional & Integrative Genomics 17: 565–581.

Cai Q, Yuan Z, Chen M, Yin C, Luo Z, Zhao X, Liang W, Hu J, Zhang D. 2014. Jasmonic acid regulates spikelet development in rice. Nature Communications 5: 3476.

Chini A, Fonseca S, Fernández G, et al. 2007. The JAZ family of repressors is the missing link in jasmonate signalling. Nature 448: 666–671.

Chung HS, Niu Y, Browse J, Howe GA. 2009. Top hits in contemporary JAZ: An update on jasmonate signaling. Phytochemistry 70: 30–40.

Egilla JN, Davies FT, Drew MC. 2001. Effect of potassium on drought resistance of hibiscus rosa-sinensis cv. Leprechaun: plant growth, leaf macro-and micronutrient content and root longevity. Plant and Soil 229: 213–224.

Elumalai RP, Nagpal P, Reed JW. 2002. A mutation in the Arabidopsis kt2/kup2 potassium transporter gene affects shoot cell expansion. Plant Cell 14: 119–131.

Feys BJF, Benedetti CE, Penfold CN, Turner JG. 1994. Arabidopsis mutants selected for resistance to the phytotoxin coronatine are male sterile, insensitive to methyl jasmonate, and resistant to a bacterial pathogen. Plant Cell 6: 751–759.

Ghosh S, Gupta SK, Jha G. 2014. Identification and functional analysis of AG1-IA specific genes of Rhizoctonia solani. Current Genetics 60: 327–341.

Gruber BD, Giehl RFH, Friedel S, von Wirén N. 2013. Plasticity of the Arabidopsis root system under nutrient deficiencies. Plant Physiology 163: 161–179.

Hakata M, Muramatsu M, Nakamura H, et al. 2017. Overexpression of TIFY genes promotes plant growth in rice through jasmonate signaling. Bioscience, Biotechnology, and Biochemistry 81: 906–913.

Hardter R. 2002. Potassium and biotic stress of plants. In: Johnston AE, ed. Feed the soil to feed the people. The role of potash in sustainable agriculture. International Potash Institute Basel, Switzerland, 345–362.

Hsu Y-Y, Kao CH. 2011. Nitric oxide is involved in methyl jasmonate-induced lateral root formation in rice. Crop Environment and Bioinformatics 8:160–7.

Kader MA, Seidel T, Golldack D, Lindberg S. 2006. Expressions of OsHKT1, OsHKT2, and OsVHA are differentially regulated under NaCl stress in salt-sensitive and salt-tolerant rice (Oryza sativa L.) cultivars. Journal of Experimental Botany 57: 4257–4268.

Kant S, Kafkafi U. 2002. Potassium and abiotic stresses in plants. In: Pasricha NS, Bansal SK, eds, Potassium for sustainable crop production. Potash Institute of India, Gurgaon, 233–251.

Kaya C, Kirnak H, Higgs D. 2006. Enhancement of growth and normal growth parameters by foliar application of potassium and phosphorus in tomato cultivars grown at high (NaCl) salinity. Journal of Plant Nutrition 24: 357–367.

Kazan K. 2015. Diverse roles of jasmonates and ethylene in abiotic stress tolerance. Trends in Plant Science 20: 219–229.

Kobayashi T, Itai RN, Senoura T, Oikawa T, Ishimaru Y, Ueda M, Nakanishi H, Nishizawa NK. 2016. Jasmonate signaling is activated in the very early stages of iron deficiency responses in rice roots. Plant Molecular Biology 91: 533–547.

Leigh RA, Wyn Jones RG. 1984. A hypothesis relating critical potassium concentrations for growth to the distribution and functions of this ion in the plant cell. New Phytologist 97: 1–13.

Li G, Wu Y, Liu G, et al. 2017. Large-scale proteomics combined with transgenic experiments demonstrates an important role of jasmonic acid in potassium deficiency response in wheat and rice. Molecular and Cellular Proteomics 16:1889–1905.

Ma TL, Wu WH, Wang Y. 2012. Transcriptome analysis of rice root responses to potassium deficiency. BMC Plant Biology 12: 161.

Mann DG, LaFayette PR, Abercrombie LL, et al. 2012. Gateway-compatible vectors for high-throughput gene functional analysis in switchgrass (Panicum virgatum L.) and other monocot species. Plant Biotechnology Journal 10: 226–236.

Marschner H. 1995. Mineral Nutrition of Higher Plants, Academic Press, London.

Mehra P, Giri J. 2016. Rice and chickpea GDPDs are preferentially influenced by low phosphate and CaGDPD1 encodes an active glycerophosphodiester phosphodiesterase enzyme. Plant Cell Report 35: 1699–1717.

Mehra P, Pandey BK, Giri J. 2017. Improvement in phosphate acquisition and utilization by a secretory purple acid phosphatase (OsPAP21b) in rice. Plant Biotechnology Journal 15: 1054–1067.

Mengel K, Viro M. 1974. Effect of Potassium Supply on the Transport of Photosynthates to the Fruits of Tomatoes (Lycopersicon esculentum). Physiologia Plantarum 30:295–300.

Murashige T, Skoog F. 1962. A Revised Medium for Rapid Growth and Bio Assays with Tobacco Tissue Cultures. Physiologia Plantarum 15: 473–497.

Oh Y, Baldwin IT, GÃlis I. 2012. NaJAZh Regulates a Subset of Defense Responses against Herbivores and Spontaneous Leaf Necrosis in Nicotiana attenuata Plants. Plant Physiology 159: 769–788.

Pandey BK, Mehra P, Verma L, Bhadouria J, Giri J. 2017. OsHAD1, a haloacid dehalogenase-like APase enhances phosphate accumulation. Plant Physiology 174: 2316–2332.

Pauwels L, Barbero GF, Geerinck J, et al. 2010. NINJA connects the co-repressor TOPLESS to jasmonate signalling. Nature 464: 788–91.

Peng X, Hu Y, Tang X, Zhou P, Deng X, Wang H, Guo Z. 2012. Constitutive expression of rice WRKY30 gene increases the endogenous jasmonic acid accumulation, PR gene expression and resistance to fungal pathogens in rice. Planta 236: 1485–1498.

Qi Z, Spalding EP. 2004. Protection of plasma membrane K+ transport by the salt overly sensitive1 Na+/H+ antiporter during salinity stress. Plant Physiology 136: 2548–2555.

Raya-González J, Pelagio-Flores R, López-Bucio J. 2012. The jasmonate receptor COI1 plays a role in jasmonate-induced lateral root formation and lateral root positioning in Arabidopsis thaliana. Journal of Plant Physiology 169: 1348–1358.

Rich CI, Black WR. 1964. Potassium exchange as affected by cation size, pH and mineral structure. Soil Science 97: 384–390.

Rich CI. 1964. Effect of cation size and ph on potassium exchange in nason soil. Soil Science 98: 100–106.

Riemann M, Dhakarey R, Hazman M, Miro B, Kohli A, Nick P. 2015. Exploring jasmonates in the hormonal network of drought and salinity responses. Frontiers in Plant Science 6: 1077.

Sarwar M. 2012. Effects of potassium fertilization on population build up of rice stem borers (lepidopteron pests) and rice (Oryza sativa l.) yield. Journal of Cereals and Oilseeds 3: 6–9.

Sawhney BL, Zelitch I. 1969. Direct determination of potassium ion accumulation in guard cells in relation to stomatal opening in light. Plant Physiology 44: 1350–1354.

Shankar A, Singh A, Kanwar P, Srivastava AK, Pandey A, Suprasanna P, Kapoor S, Pandey GK. 2013. Gene expression analysis of rice seedling under potassium deprivation reveals major changes in metabolism and signaling components. PLoS One 8: e70321.

Sheard LB, Tan X, Mao H, et al. 2010. Jasmonate perception by inositol-phosphate-potentiated COI1-JAZ co-receptor. Nature 468: 400–405.

Singh AP, Pandey BK, Deveshwar P, Narnoliya L, Parida S.K, Giri J. 2015. JAZ Repressors: Potential involvement in nutrients deficiency response in rice and chickpea. Frontiers in Plant Science 6: 975.

Sparks DL. 1987. Potassium dynamics in soils. In: Stewart BA, ed. Advances in soil science. Springer, New York, 1–63.

Staswick PE, Su W, Howell SH. 1992. Methyl jasmonate inhibition of root growth and induction of a leaf protein are decreased in an Arabidopsis thaliana mutant. Proceedings of the National Academy of Sciences 89: 6837–6840.

Takehisa H, Sato Y, Antonio BA, Nagamura Y. 2013. Global transcriptome profile of rice root in response to essential macronutrient deficiency. Plant signaling and behavior 8: e24409.

Terry N, Ulrich A. 1973. Effects of potassium deficiency on the photosynthesis and respiration of leaves of sugar beet. Plant Physiology 51: 783–786.

Tester M, Blatt MR. 1989. Direct measurement of k+ channels in thylakoid membranes by incorporation of vesicles into planar lipid bilayers. Plant Physiology 91: 249–252.

Thatcher LF, Cevik V, Grant M, Zhai B, Jones JD, Manners JM, Kazan K. 2016. Characterization of a JAZ7 activation-tagged Arabidopsis mutant with increased susceptibility to the fungal pathogen Fusarium oxysporum. Journal of Experimental Botany 67: 2367–2386.

Thines B, Katsir L, Melotto M, Niu Y, Mandaokar A, Liu G, Nomura K, He SY, Howe GA, Browse J. 2007. JAZ repressor proteins are targets of the SCF(COI1) complex during jasmonate signalling. Nature 448: 661–665.

Troufflard S, Mullen W, Larson TR, Graham IA, Crozier A, Amtmann A, Armengaud P. 2010. Potassium deficiency induces the biosynthesis of oxylipins and glucosinolates in Arabidopsis thaliana. BMC Plant Biology 10: 172.

Wang M, Zheng Q, Shen Q, Guo S. 2013. The critical role of potassium in plant stress response. International Journal of Molecular Sciences 14: 7370–7390.

Wasternack C, Hause B. 2013. Jasmonates: Biosynthesis, perception, signal transduction and action in plant stress response, growth and development. An update to the 2007 review in annals of botany. Annals of Botany 111: 1021–1058.

Wasternack C, Song S. 2017. Jasmonates: biosynthesis, metabolism, and signaling by proteins activating and repressing transcription. Journal of Experimental Botany 68: 1303–1321.

Wu H, Ye H, Yao R, Zhang T, Xiong L. 2015. OsJAZ9 acts as a transcriptional regulator in jasmonate signaling and modulates salt stress tolerance in rice. Plant Science 232: 112.

Wu W, Peters J, Berkowitz GA. 1991. Surface charge-mediated effects of Mg^2^+ on K+ flux across the chloroplast envelope are associated with regulation of stromal pH and photosynthesis. Plant Physiology 97: 580–587.

Yamada S, Kano A, Tamaoki D, Miyamoto A, Shishido H, Miyoshi S, Taniguchi S, Akimitsu K, Gomi K. 2012. Involvement of OsJAZ8 in jasmonate-induced resistance to bacterial blight in rice. Plant Cell Physiology 53: 2060–72.

Yang DL, Yao J, Mei CS, et al. 2012. Plant hormone jasmonate prioritizes defense over growth by interfering with gibberellin signaling cascade. Proceedings of the National Academy of Sciences 109: E1192–1200.

Ye H, Du H, Tang N, Li X, Xiong L. 2009. Identification and expression profiling analysis of TIFY family genes involved in stress and phytohormone responses in rice. Plant Molecular Biology 71: 291–305.

